# Cultivating *in situ* triggers growth of previously uncultivated microorganisms via a growth initiation factor in nature

**DOI:** 10.1101/590653

**Authors:** Dawoon Jung, Koshi Machida, Yoichi Nakao, Tomonori Kindaichi, Akiyoshi Ohashi, Yoshiteru Aoi

## Abstract

Most microorganisms resist cultivation under standard laboratory conditions. On the other hand, to cultivate microbes in a membrane-bound device incubated in nature (*in situ* cultivation) is an effective approach. In the present study, we applied *in situ* cultivation to isolate diverse previously uncultivated marine sponge-associated microbes and comparatively analyzed this method’s efficiencies with those of the conventional method. Then, we attempted to clarify the key and unknown mechanism of *in situ* cultivation by focusing on growth triggering via growth initiation factor. We hypothesized that majority of environmental microorganisms are in nongrowing state and requiring “growth initiation factor” for the recovery and that can be provided from environments. Consequently, significantly more novel and diverse microbial types were isolated via *in situ* cultivation than by standard direct plating (SDP). Next, the effect of the sponge extract on starvation recovery was compared between strains derived from *in situ* and SDP cultivation. Adding small amounts of the sponge extracts to the medium elevated the colony-formation efficiencies of the *in situ* strains at the starvation recovery step, while it showed no positive effect on that of SDP strains. Conversely, specific growth rates or carrying capacities of all tested strains were not positively affected. These results indicate that, 1) the sponge extract contains chemical compounds that facilitate starvation recovery, these substances selectively worked on the *in situ* strains, and 2) growth initiation factor in the sponge extract did not continuously promote growth activity but worked as triggers for regrowth (resuscitation from dormancy).

**Importance:** Most microbial species resist cultivation under laboratory condition. This is critical impediment for both academic and applied microbiology, and thus clarification of the mechanism of microbial uncultivability is highly demanded. Several evidences have been reported that to cultivate microbes in a membrane-bound device incubated in nature (*in situ* cultivation) is an effective approach. However, the mechanism behind this approach has not been clarified. The present study shows the evidence that 1) initiating growth is a key for cultivating previously uncultivated microbes rather than simple growth promotion, and 2) growth initiation factor (signaling-like compounds) in natural environments stimulate microbial resuscitation from a nongrowing state. Since no study has focused on growth initiation for cultivation of previously uncultivated microorganisms, the discovery shown in the present study provides a new insight into microorganisms previously considered uncultivable and a microbial growth controlling system in nature.

## Introduction

Most microbes remain uncultivated and have been referred to as ‘mysterious dark matter’ of the microbial world (1, 2). Cultivation independent survey have demonstrated great diversity among these uncultivated species (3–5). Accessing this “missing” microbial diversity is of great interest to both basic and applied sciences and has been regarded as a major challenge for microbiology.

Standard agar plating as a conventional method of cultivating microorganisms is limited because a significantly low proportion (usually less than 1%) of the plated microbes readily form visible colonies on the agar plates, thus leading to plate count anomalies (6, 7). To overcome the limitations, much effort has been devoted to developing alternative approaches, including physically separating cells to decrease competition or inhibitors (8–10), using modification to prepare agar media, using alternative gelling agents or antioxidants to minimize unfavorable compounds (11–13), and adding signal molecules or cocultivating with recruiter organisms to better reflect the natural environment (14–16). Nevertheless, most postulated extant microbes in nature remain uncultivated; thus, other essential factors for microbial cultivation that exist in nature are likely absent from those artificial conditions.

One simple solution for approaching the unknown growth factors is to incubate the microbes in their natural environment. Applying this idea to microbial cultivation led to developing *in situ* cultivation methods aiming to better simulate the natural environment (for a review, see Epstein et al., 2010 [17]). Several *in situ* cultivation methods with similar basic concepts have been applied to various environments, including sediment, activated sludge, alkaline soda lakes, sponges, soil, and a hot spring environment, and have been demonstrated to be highly capable of microbial cultivation (18–24). However, why these *in situ* cultivation techniques enable cultivating previously uncultivated microbes that are otherwise difficult or impossible to grow using conventional methods remains unclear. Answering this question would provide a key factor for microbial cultivation and contribute to developing additional advanced cultivation techniques.

In the present work, we applied *in situ* cultivation to the marine sponge to isolate previously uncultivated microbial species. Marine sponges are rich sources of bioactive secondary metabolites of biotechnological interest for their antiviral, antitumoral, antimicrobial and cytotoxic properties (25–27). It has been suggested that symbiotic microbes in marine sponges produces some of these bioactive metabolites (28–32). Additionally, recently developed culture-independent approaches have provided molecular evidence of these microorganisms’ functional roles (33–35). However, most of these producers remain unexplored and unavailable because the microbes associated with the sponges cannot be easily cultivated (32).

One focus of the present study was to determine an effective cultivation method to broaden the accessible marine sponge-associated microbes. To do this, we employed one of *in situ* cultivation method, diffusion chamber to isolate previously uncultivated marine sponge-associated microbes and comparatively analyzed this method’s efficiencies with those of the conventional method (direct agar plating). Although the applicability of *in situ* cultivation to marine (22) and freshwater sponges (21) has been demonstrated in previous studies, this approach’s potential for cultivating novel, previously uncultivated microbial species from sponge samples remains unknown.

Another aim of the study is clarifying the reason why *in situ* cultivation enables growing previously uncultivated microbes that cannot be isolated by conventional methods. We hypothesized that inactive or nongrowing microbial cells are stimulated and aroused to begin regrowing during *in situ* incubation by a growth initiation factor from the outside environment and enriched inside an *in situ* cultivation chamber with nutrients. This brings different microbial groups, which mostly resist cultivation by conventional methods, to the culture collection. To verify our hypothesis, recovery from the nongrowing state in response to the marine sponge extract was comparatively analyzed between microbes isolated via *in situ* cultivation and those isolated via the conventional method.

## Results

### Identifying isolates based on the 16S rRNA gene

Figures 1 presents the overall experimental design for the cultivation experiments (Fig. 1a), structure and application of the device (DC) (Fig. 1b and c), and principle of *in situ* cultivation method (Fig. 1d). One hundred twenty bacterial isolates (60 per cultivation method) were randomly selected and identified. *In situ* cultivation (DC) enabled isolating 37 species (defined as operational taxonomic units [OTUs] composed of 16S rRNA gene sequences sharing over 97% identity) from six taxonomic groups (*Actinobacteria*, *Bacteroidetes*, *Firmicutes*, *Alphaproteobacteria*, *Epsilonproteobacteria* and *Gammaproteobacteria*) (Fig. 2, Table S1). Standard direct plating (SDP) cultivation enabled isolating 13 species from three taxonomic groups (*Bacteroidetes*, *Alphaproteobacteria* and *Gammaproteobacteria*) (Table S2). These results suggested that *in situ* cultivation (DC) yielded significantly higher diversity among the isolates at the species level than among those obtained via SDP cultivation (Fig. 2). In addition, a few species overlapped among the isolates from each method (Fig. 2). One species belonging to *Ruegeria atlantica* (99% similarity to the closest known species) was shared between the DC and SDP isolates.

**Figure 1.**
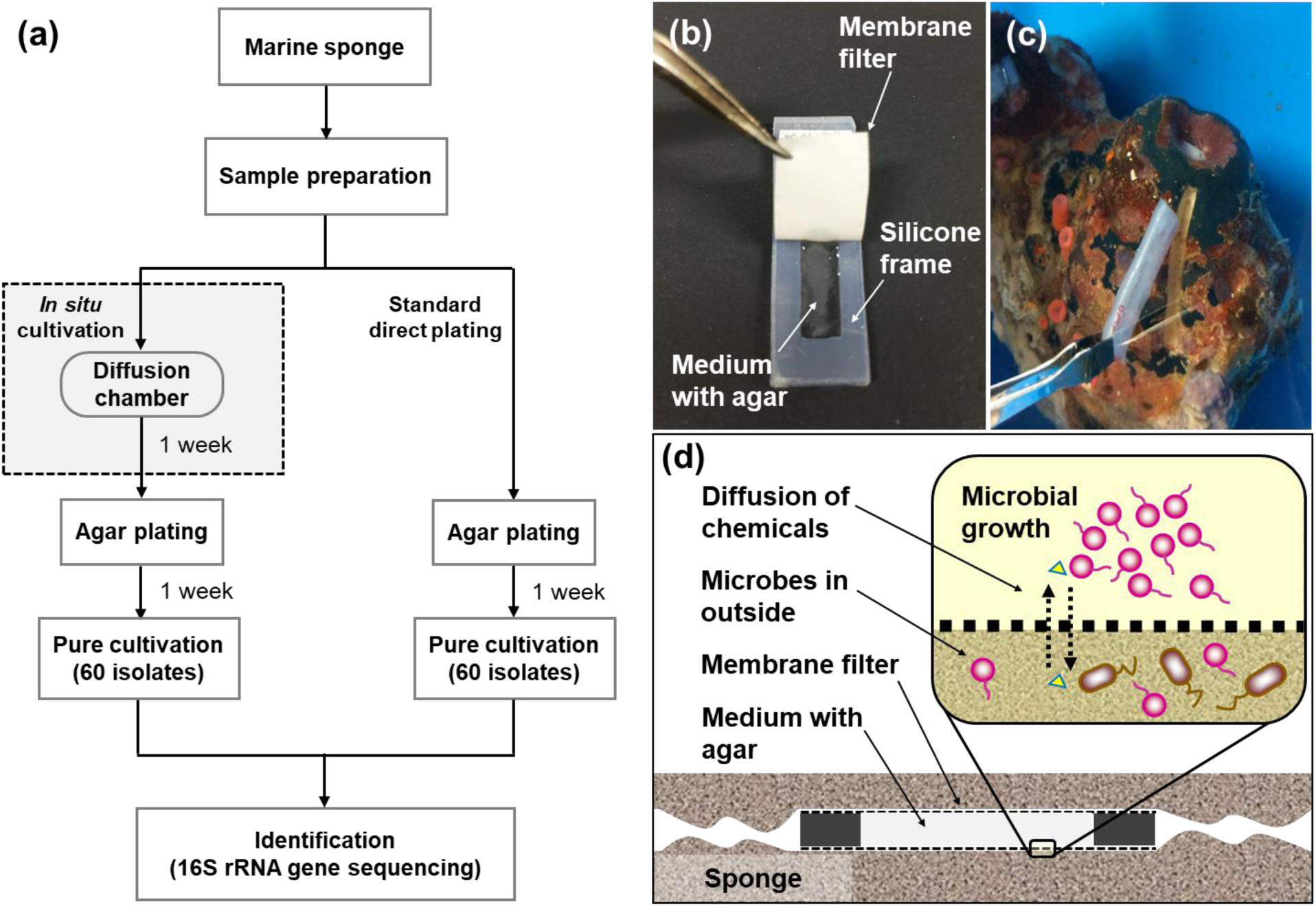
Flowchart of the experimental steps for microbial cultivation from marine sponges (a) and diffusion chamber (DC) cultivation method applied in the present study: a photo showing structure of the chamber (b), a photo showing installation of DC device into the marine sponge (c), a schematic image showing the principle of DC (d).

**Figure 2.**
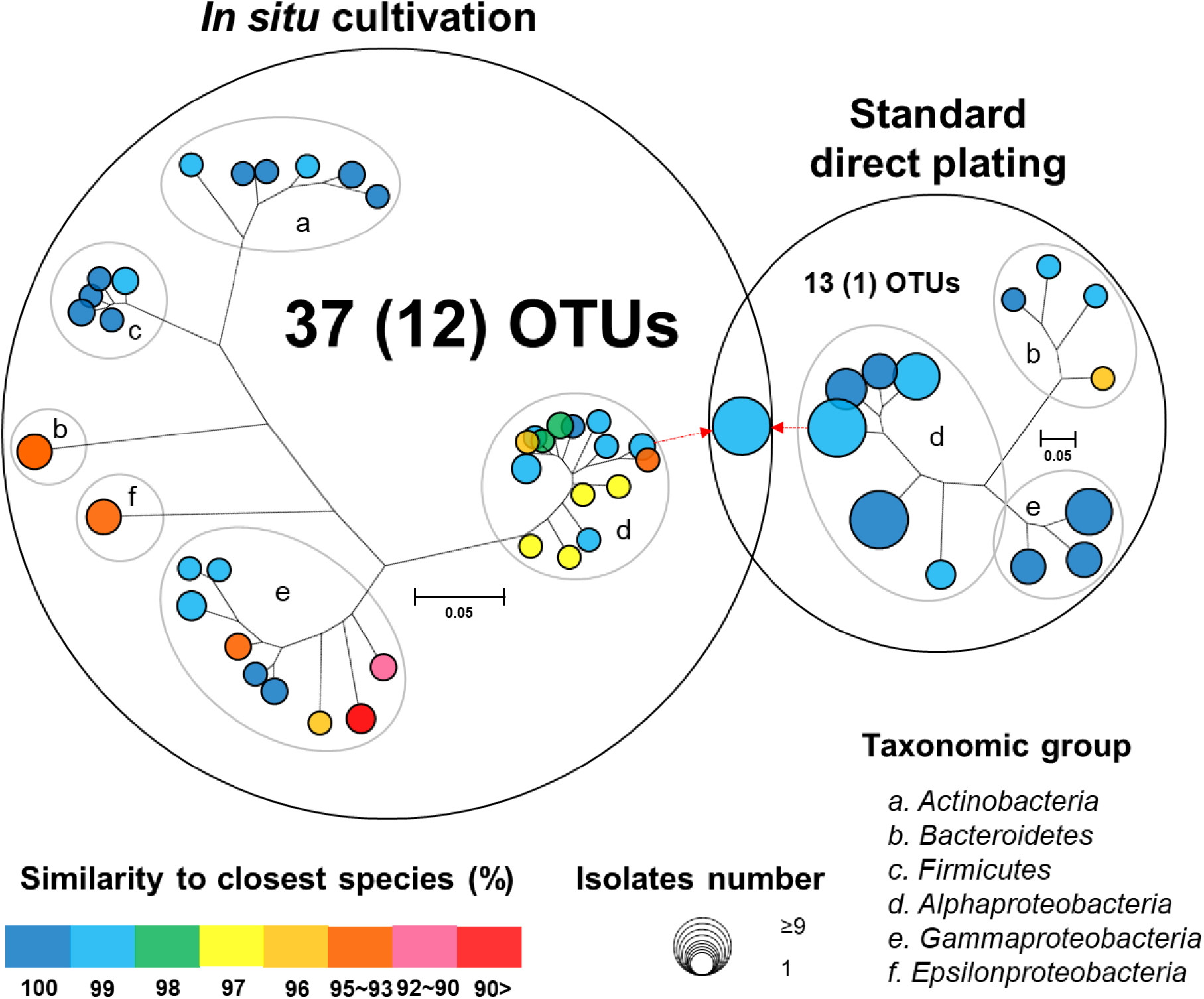
Venn diagram consisting of phylogenetic trees based on the 16S rRNA gene of isolates from each cultivation method. The trees are maximum-likelihood trees (fast bootstrap, 1,000 replicates). Circle size and color represent the number of OTUs (defined at 97% 16S rRNA gene sequence identity) and 16S rRNA similarity to the closest known relative in GenBank, respectively. Represented isolates are also grouped into taxonomic groups (a to f) at the phylum level (class level for Alpha, Gamma and Epsilon Proteobacteria) by gray elliptical circles. Circles with pointed arrows indicate species that are overlapped among the isolates in the different cultivation methods. The numbers in parentheses show the numbers of novel species. The outer circle area corresponds to the number of isolated OTUs from each cultivation method.

The ratios of novel species, defined as a strain with ≤97% 16S rRNA similarity to the closest known relative among the isolates, differed between the SDP and *in situ* cultivation methods. Only one SDP isolate (1.7%) was a novel species, while 40% (24 isolates belonging to 12 species) of the *in situ* isolates was novel species (Fig. 2, 3).

### Microbial community compositions among the samples: comparison of the culture-dependent and -independent methods

The microbial community compositions of the marine sponge (*Theonella swinhoei*) samples were analyzed by Illumine-MiSeq sequencing based on the 16S rRNA gene. A total of 114,624 reads with a median length of 252 base pairs (bp) (V4∼533–786 bp) assigned to 1625 OTUs were obtained from the sample. Ten major taxonomic groups were detected (mostly at the phylum level but at the class level for Proteobacteria): *Acidobacteria*, *Actinobacteria*, *Bacteroidetes*, *Chloroflexi*, *Gemmatimonadetes*, *Poribacteria*, *Alphaproteobacteria*, *Epsilonproteobacteria*, *Gammaproteobacteria* and SBR1093 (with a cut-off of <1.0%; Fig. 4). The *in situ* isolates belonged to six groups, *Actinobacteria*, *Bacteroidetes*, *Firmicutes*, *Alphaproteobacteria*, *Epsilonproteobacteria* and *Gammaproteobacteria*, while the SDP isolates belonged to three groups: *Bacteroidetes*, *Alphaproteobacteria* and *Gammaproteobacteria*.

**Figure 3.**
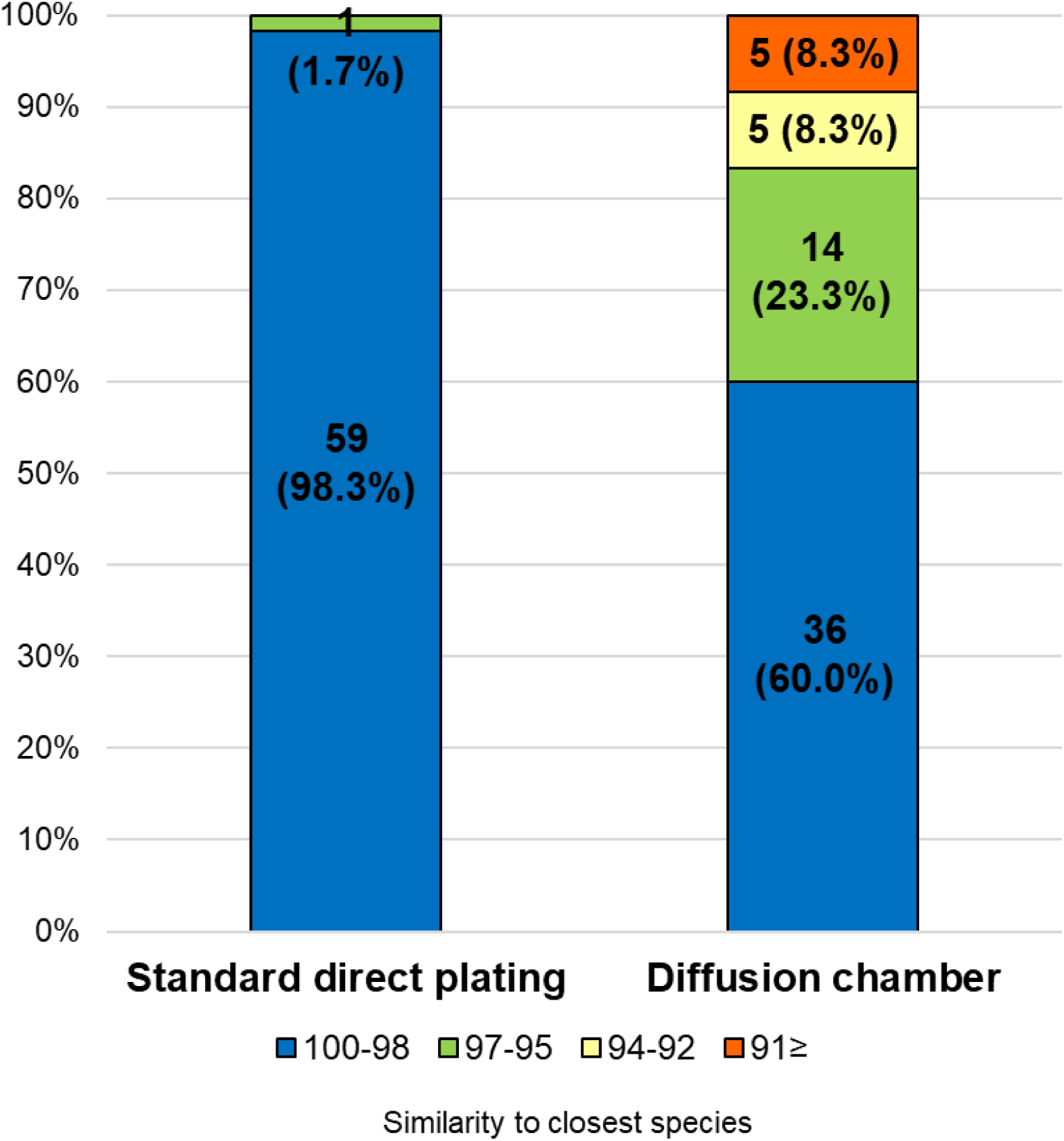
Similarity among the isolates from each cultivation method based on the 16S rRNA gene to the closest known relative in GenBank. Numbers in the bar graphs represent OTUs for each similarity level and its ratio (%).

**Figure 4.**
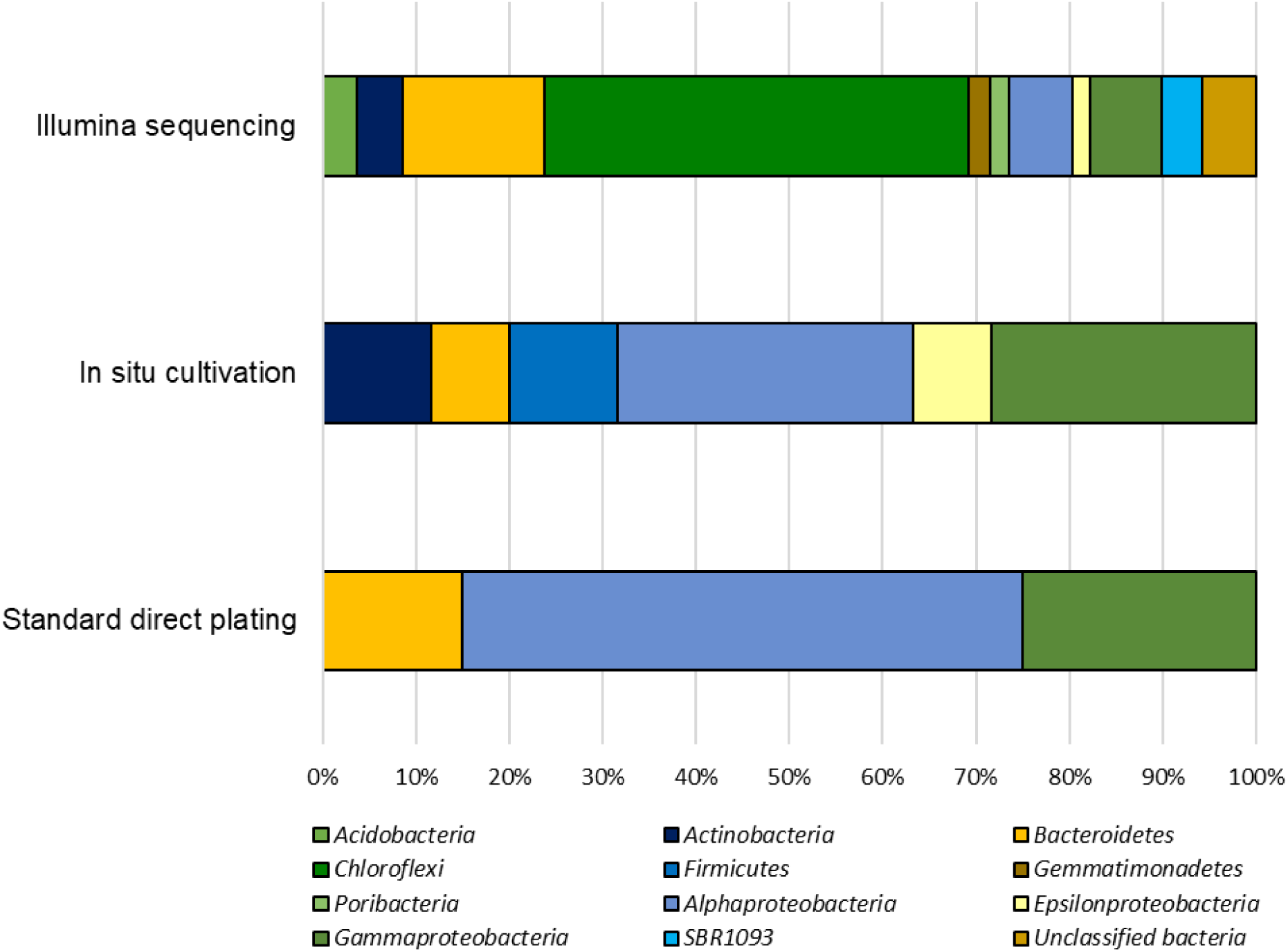
Phylogenetic distribution of 16S rRNA gene sequences of sponge-associated bacteria from the original sample (*Theonella swinhoei*) and isolates obtained by *in situ* cultivation and standard direct plating methods.

### Effect of the sponge extract on starvation recovery

For further investigation, we picked 28 and 11 strains obtained from *in situ* and SDP cultivation, respectively, which are representing identical species of all identified isolates except a few lost species (we hereafter define those selected stains as *in situ* and SDP strains, respectively). Figure 5a shows the effect of adding the sponge extract (0.1% of the total volume) to the medium on starvation recovery (colony formation efficiency). Each strain’s colony efficiency ratio between the two culture conditions (with and without the sponge extract) was calculated. Adding the sponge extract to the medium exerted different effects on the recoveries between the *in situ* and SDP strains. Adding the sponge extract positively affected the recoveries of all tested *in situ* strains. Among them, 11 strains (11/28, 39%) more than doubled in colony numbers on the agar media with the sponge extract compared with that on the media with no sponge extract. In contrast, none of SDP strains was not positively affected or rather negatively affected on the recovery by the sponge extract addition, except two strains were slightly positively affected (Fig. 5a). Note that closest species of one SDP strain (SDP8) showed positive is *Ruegeria atlantica*, which is commonly found in *in situ* isolates (Table S1 and S2).

**Figure 5.**
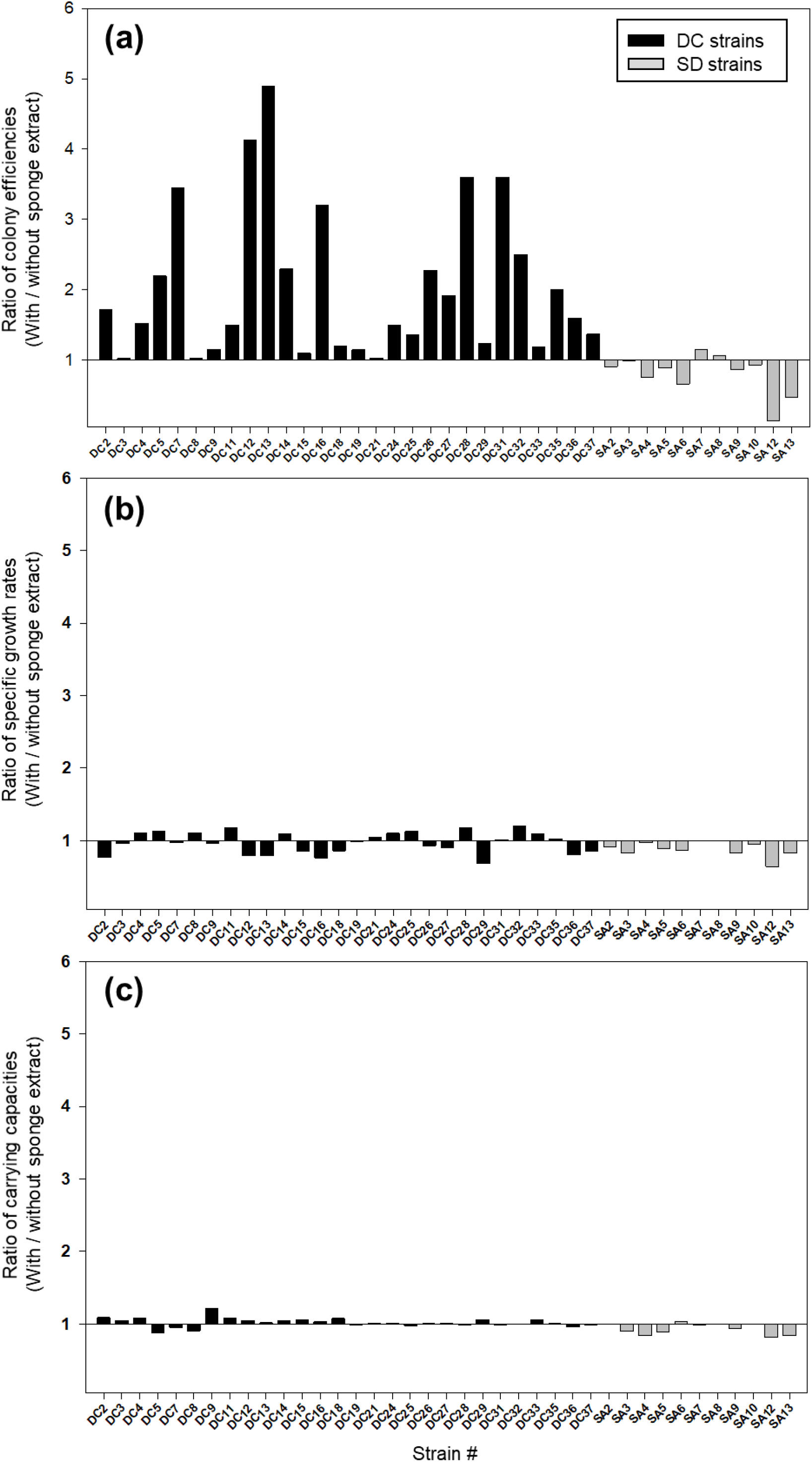
Effect of the sponge extract on starvation recovery (a), specific growth rate (b) and carrying capacity (c) of the *in situ* and SDP strains. The ratio of colony efficiencies between the value measured under two culture conditions (colony numbers on the agar medium with the sponge extract/colony numbers on the agar medium without the sponge extract) was calculated for each selected strain (a). The ratio of the specific growth rate (*μ*) between the value measured under the two culture conditions (*μ*) with the sponge extract/*μ* without the sponge extract) was calculated for each tested strain (b). The ratio of the carrying capacity (maximum value of OD600) between the value measured under the two culture conditions (carrying capacity with the sponge extract/ carrying capacity without the sponge extract) was calculated for each tested strain (c).

### Effect of the sponge extract on specific growth rates and carrying capacities

Figure 5b and c shows the effect of adding the sponge extract (0.1% of the total volume) to the medium on the specific growth rate and carrying capacity, respectively. Each strain’s specific growth rate and carrying capacity were measured and compared under two medium conditions (with and without the sponge extract) to examine the sponge extract’s effect on growth activity (Fig. S1). In contrast to its effect on starvation recovery (colony forming efficiencies), adding the sponge extract to the medium did not significantly elevate or decrease both the specific growth rate and carrying capacity of any tested strain. The addition slightly negatively affected the specific growth rates for more than half the tested *in situ* strains (15/28, 54%) and all SDP strains except SDP7 and SDP8 (SDP8 is commonly isolated species from *in situ* cultivation).

## Discussion

### Effectiveness of *in situ* cultivation techniques for isolating novel species from marine sponges

Marine sponges are sources of many bioactive natural products (36–38), which are often produced by host-specific microbes that are mostly unknown because of their uncultivability (32, 39). Molecular surveys have shown that sponges host rich microbial communities (40, 41); however, only a minor component of this richness has been cultivated via the conventional cultivation method (32). Extensive cultivation efforts have made use of several alternative techniques to isolate previously uncultured sponge-associated bacteria, including adding the sponge extracts or its skeleton to the media (42, 43), using oligotrophic media (36), adding antibiotics to inhibit fast-growing bacteria (44), using alternative gelling agent (45), and using *in situ* diffusion devices (21, 22) and floating filters (42). These alternative approaches yielded an increased novelty of sponge isolates and improved cultivability rates up to 14% in some cases (42). However, most postulated extant microbes in sponges remain uncultivated; thus, further efforts to discover such novel microbes are needed. Here, we isolated previously uncultivated microorganisms from marine sponges more efficiently than in previous studies (40% of *in situ* isolates were novel species in this study). Newly discovered microbes are candidates as sources for valuable secondary metabolites, and the newly established approach would be a strong tool for further accessing untapped microbial resources.

### Hypothesis: a key mechanism of *in situ* cultivation for growing previously uncultivated microbial types

*In situ* cultivation approach via diffusion chamber (DC) enabled obtaining a different culture collection that was larger and more novel than that obtained by standard direct plating (SDP) (Fig. 2, 3). Only one species was common in culture collections between the *in situ* and SDP cultivation methods. These results indicated that most isolated species were unique to their isolation approach, and *in situ* cultivation enabled cultivating specific microbial types that have rarely been isolated by conventional cultivation approaches.

These results raised the question, “what mechanism from the alternative approach yielded these previously uncultivated microbial groups that differed entirely from the standard cultivation approach?”. A simple explanation is that during *in situ* cultivation, inoculated microbes receive unknown growth components from the natural environment necessary for their growth but absent from the artificial medium (46). However, the present study found that isolates unique to *in situ* cultivation grew stably under the same conditions as those of the SDP cultivation at the subcultivation step, which lacks such growth components. This phenomenon also occurred in previous studies that used *in situ* cultivation (18, 20, 21, 24). The only the difference between the *in situ* and SDP cultivation methods in the experimental procedure was whether a one-week incubation was performed in the natural environment (inside the marine sponge in this study) prior to agar plate cultivation (Fig. 1, a). We then explored what mechanism could explain this observation.

We hypothesized that 1) most environmental microbes (in this case, microbes associated with marine sponges) are in nongrowing states (viable but nonculturable [VBNC], dormancy, near-zero growth, inactive state) to survive under nutrient-limited conditions (47), and 2) they require a specific signaling-like compounds, so called growth initiation factor to recover from the nongrowing state and begin regrowth together with nutrient (here, it is supplied by medium in diffusion chamber). Such growth initiation factor exists in natural environments but is usually absent from artificial media. We expected that this factor stimulated and aroused inactive microbial cells (in a nongrowing state), resulting in the growth of specific microbial groups that could not be isolated via standard cultivation (Fig. 6). However, although considered essential for microbial cultivation, the microbial awakening mechanism remains unclear (48).

**Figure 6.**
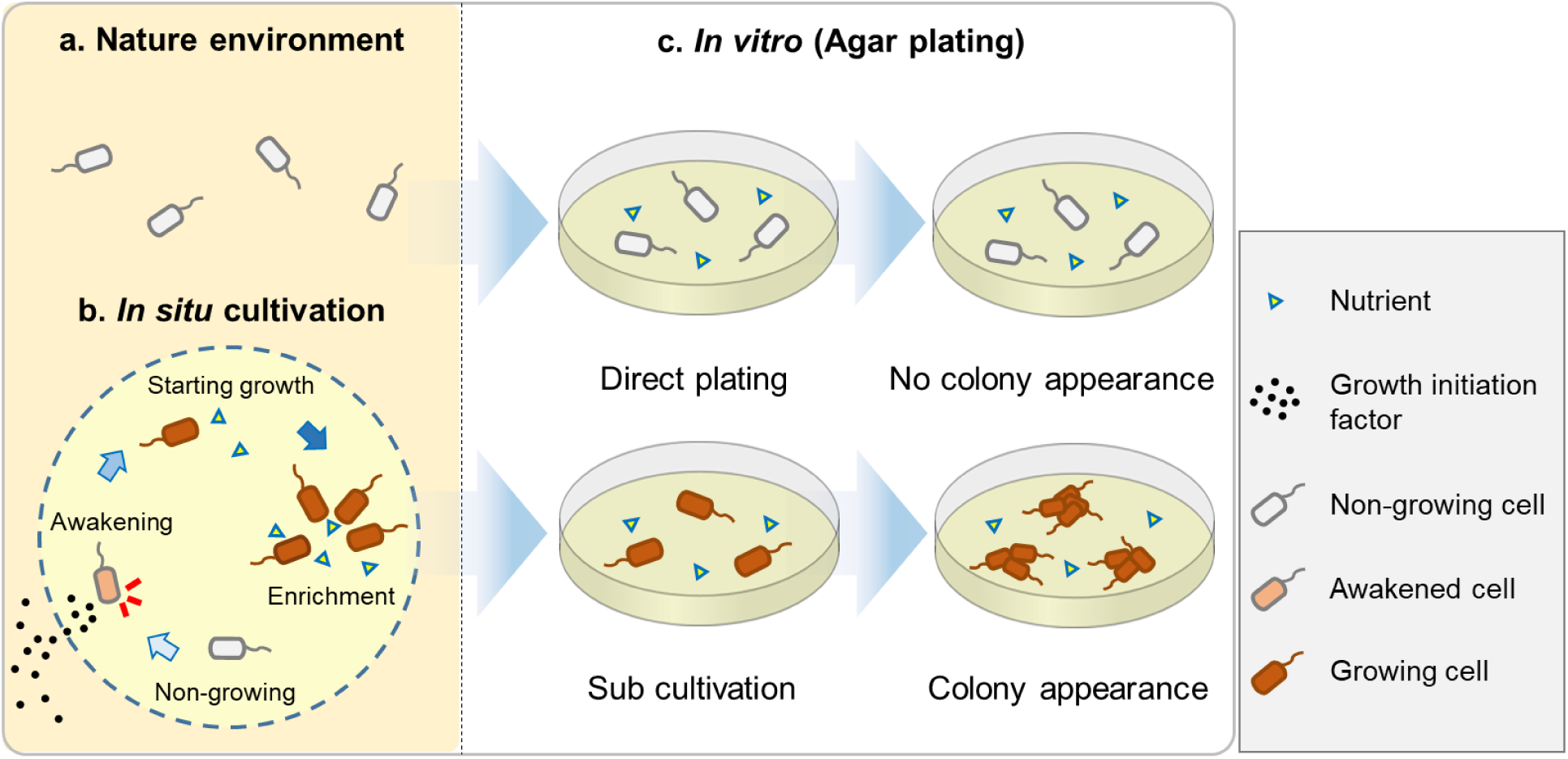
Hypothesis explaining how *in situ* cultivation yields different culture collections from microbial recovery in a natural environment; certain microbial types are non-growing (dormant) to survive unfavorable conditions such as nutrient limited condition (a); during *in situ* cultivation dormant microbes resuscitate from non-growing state stimulated by growth initiation factor provided from outside environments, then they started to grow and enriched inside supported by sufficient nutrients in the chamber (b); non-growing microbes requiring growth initiation factor for their awakening, do not start to grow when they were directly inoculated on an agar plate, and resulting no visible colony appearance; on the other hand, when they were sub-cultured after the *in situ* cultivation, they continuously grow on an agar plate, and resulting visible colony appearance (c).

### Growth triggering by growth initiation factor in nature

To verify the hypothesis and clarify the mechanism behind the *in situ* cultivation results, the effect of adding small amounts of the sponge extract on the starvation recoveries and specific growth rates of representative isolates (from every species) were compared between the strains derived from *in situ* and SDP cultivation. Most species (represented isolates) from the *in situ* and SDP cultivations were used for the test, except a few strains which had been lost.

The results supported the hypothesis as follows. Adding the sponge extract to the medium (only 0.1% of the total volume) elevated the colony formation efficiencies of all tested *in situ* strains at the starvation recovery step, while it only negatively affected that of the SDP strains (Fig. 5a). In contrast, adding the sponge extract did not elevate the specific growth rates in either the *in situ* or SDP strains but rather slightly suppressed them, especially the SDP strains (Fig. 5b). In addition, adding the sponge extract did not positively affect the carrying capacity of every tested strain (Fig. 5c). These results show that 1) the sponge extract contains a growth initiation factor that facilitate starvation recovery, and these substances selectively worked on the *in situ* strains, and 2) the growth initiation factor in the sponge extract did not continuously promote growth activity but worked as triggers for regrowth, which was likely resuscitation from dormancy (nongrowing state).

Therefore, during the *in situ* cultivation, such signaling-like compounds, defined as “growth initiation factor”, was provided from the natural environment (i.e., the marine sponge), while nutrients were provided inside the chamber from the beginning. These factors enabled microbes begin to growing again and finally become enriched inside the chamber (Fig. 6b). In contrast, standard agar plate medium lacks such growth initiation factor; thus, significant numbers of inoculated microbial cells did not resuscitate from dormancy and thus did not form colonies (Fig. 6c, upper part). Because sponge-associated microbes produce various chemical compounds, such growth initiation factor are likely also produced by specific microbes associated with the sponge (32).

In contrast, because SDP strains require no growth initiation factor to begin regrowing, these strains are frequently isolated via conventional approaches (the SDP method in the present study) and have been discovered in previous studies. Therefore, microbial types in isolates differ completely between methods, and the novelty and diversity of isolates derived from *in situ* cultivation are higher than are those derived from the conventional approach.

Additionally, the sponge extract was highly toxic to the microbes. Adding the sponge extract to the media at higher concentrations than that of the original test condition (e.g., 1%, which was 10 times higher than the original test condition) strongly inhibited isolate growth (data not shown). This was likely due to antimicrobial substances in the sponge, such as secondary antimetabolites or high concentrations of toxic trace elements, which can accumulate in the sponge body (43, 49, 50). Furthermore, SDP strains are more sensitive to that toxicity than are *in situ* strains as described above and shown in Fig. 5. In other words, under the tested condition (0.1%), the negative effect of adding the sponge extract appeared only for SDP strains. This suggests that microbial types that are susceptible to being enriched by *in situ* cultivation are less sensitive to the toxic substances that are likely to be excreted from the sponge in nature; thus, microbial types derived from *in situ* cultivation would be more competitive in the original environment (inside the sponge). We reported a similar observation in our previous study, suggesting that *in situ* isolates are more competitive under their original conditions (natural environment) than are SDP isolates (23). Therefore, *in situ* isolates might outcompete the microbes that are frequently isolated via SDP cultivation. This might be another reason why *in situ* isolates are novel and differ phylogenetically from SDP isolates.

### Significance of the present study relative to solving microbial uncultivability

Various *in situ* cultivation methods with similar basic rationales have been applied to various environmental samples and have been shown to be highly capable of microbial cultivation compared with conventional approaches because the method led to isolating some uncultivated and phylogenetically new and industrially important isolates (18-21, 51–53). Furthermore, mining “microbial dark matter” using this new approach remains highly expected (2). However, no study has clarified the growth facilitation mechanism of *in situ* cultivation or the reason why *in situ* cultivation yields more phylogenetically novel and diverse microbial types than does the conventional method. This has limited further advancing the method regardless of its potential. The present study resolved this issue for the first time.

Furthermore, the discovery in the present study, especially regarding microbial resuscitation from a nongrowing state in response to the growth initiation factor gives us an important clue toward solving this challenging issue of why most environmental microbes cannot grow on standard media. This has been described as “the great plate count anomaly” (6, 7). Although most microbial types that resist cultivation are assumed to be fastidious and require culture conditions within a narrow range, the results of the present study suggest that microbial uncultivability cannot simply be explained by the unfitness of the specific strain to certain culture conditions such as medium composition, gas composition, temperature, and pH (54).

Several studies have been conducted to determine the reasons for microbial uncultivability, such as inoculum size (55) and phosphate-catalyzed hydrogen peroxide formation from agar (12, 13, 56). A few attempts have been made to discover microbial interactions via signaling molecules that promote the growth of specific microbial types. Short peptides (57), siderophores (58) and quinones (59) have been identified as growth-promoting factors for microbial types that cannot grow independently.

However, the causes of the uncultivability and the mechanism facilitating growth of previously uncultivated microbes shown in the present study differ from those reported in previous studies in several aspects as follows. First, no study has focused on growth triggering or distinguished this from simple growth promoting, yet that must be the key to solving the microbial uncultivability as shown herein. Second, no study has provided experimental evidence showing that the growth of diverse environmental microbes is controlled by a growth initiation factor.

Previous studies have reported various types of microbial nongrowing states such as viable but nonculturable (VBNC), near-zero growth (NZG) and dormancy (60–62), which are suspected to cause microbial uncultivability. Wide species distributions in the VBNC state have been detected in broad environments such as sediment (63), estuarine water (64), seawater (65) and soil (66). The broad distribution of inactive microbial cells in environments reinforce that this state is a common mechanism for coping with unfavorable environmental conditions (60, 62). Although microbial resuscitation from dormancy remains unclear, a few studies on resuscitation or germination of gram-positive bacteria via signaling compounds have been reported (67–69). Therefore, many environmental microorganisms may possess similar systems for microbial awakening from nongrowing states via signaling compounds. Furthermore, the “scout hypothesis” theory has been proposed, which states that 1) dormant microbes stochastically and spontaneously awaken, and 2) awakened cells trigger the awakening of other dormant cells (70, 71). However, no study has experimentally confirmed that growth triggering of environmental microbes from a nongrowing state is a key phenomenon of microbial cultivation.

### Concluding remarks and perspectives

We demonstrated that *in situ* cultivation effectively isolates previously uncultivated microbial types from the marine sponge. We further clarified that *in situ* cultivation facilitates isolating novel and different microorganisms compared with the conventional approach because of the discovery of the growth-triggering phenomenon from a nongrowing state by growth initiation factor in the natural environment. To our knowledge, the present study shows the first evidence that 1) triggering growth is a key for cultivating previously uncultivated microbes, and 2) the growth initiation factor (signaling-like compounds) in natural environments (likely produced by microbes) stimulate microbial resuscitation from a nongrowing state.

Next challenge beyond the present study include 1) identifying the growth initiation factor via natural chemical approaches and 2) clarifying the donors and accepters of signaling compounds (i.e., who produces what and to whom). This will provide a new insight into microorganisms previously considered uncultivable and the complex networking system for controlling growth among microbes in nature.

## Materials and Methods

### Sample collection

Marine sponge (*Theonella swinhoei*) specimens were collected while scuba diving at 15-20 m depth by hand in August 2015, near Okino island, Kochi prefecture, Japan. Sponges were kept in a cooling box with seawater and transported to the aquarium and laboratory for further experiments.

### Media

For the microbial cultivation experiments, we used the following four media: 1) 1:10 diluted Reasoner’s 2A (R2A) media (10% of the manufacturer’s suggested concentration, Nihon Seiyaku, Japan); 2) marine media (Difco, Franklin Lakes, NJ, USA); 3) fish extract media (0.2 g fish extract and 0.1 g yeast extract per liter); and 4) sponge extract media (0.01 g peptone and 40 ml aqueous sponge extract per liter). The sponge extract was prepared by mixing homogenized sponge and sterile distilled water including 3.5% artificial sea salt at a 1:1 (vol/vol) ratio, then vortexing for 1 min, spinning down (10 min, 8,000 rpm), and filter sterilization using a 0.2-μm pore-size filter. All media contained 1.5% agar to produce solid medium, and all media except the marine media were supplemented with 3.5% artificial sea salt, SEALIFE (Marine Tech, Tokyo, Japan). Colonies grown on these media were subcultured for purification on 1.5% agar plates supplemented with 1:10 R2A broth. All isolates grew well on 1:10 diluted R2A agar media. Some isolates were derived from different media at the subcultivation or direct plating steps.

### Diffusion chamber *in situ* cultivation

Diffusion chamber (DC) i*n situ* cultivation method was performed to isolate marine-sponge associated microbes. The DC was prepared as described previously (18). The DC is an *in situ* cultivation device that allows exchanging chemical compounds, thereby simulating a natural environment (Fig. 1. d). To produce a chamber, a 0.1-μm pore-size filter polycarbonate membrane (Millipore, Darmstadt, Germany) was glued to a rectangular-shaped silicone flame (w = 3 cm × 1.2 cm, d = 0.3 cm) with a waterproof adhesive (Cemedain, Tokyo, Japan) (Fig. 1. b). To prepare the marine sponge sample inoculum, 20 g of the subsample was rinsed three times with sterile artificial sea water and pulverized using a sterile mortar and pestle with the same volume of sterile artificial sea water. The aliquots were diluted 10^-4^ to 10^-5^ with sterilized water including 3.5% artificial sea salt and mixed with warm (45°C) agar with 1:10 diluted R2A. The mix was placed on the chamber membrane, and the second membrane was glued to the other side of the silicone flame, sealing the agar inside to form a chamber.

For *in situ* cultivation, the sponges were kept in aquariums in the Takehara station of Hiroshima University in Takehara-city, Japan. The aquarium was a cylindrical container with a holding capacity of ca. 43.3 L (Ø = 0.35 m, d = 0.45 m), and seawater was supplied continuously. To install the DC into the sponge, similar sized grooves as those of the device were made on the specimen’s surface by cutting with a scalpel blade, and four DCs were inserted (Fig. 1. c). After one week of *in situ* incubation, DCs were retrieved from the sponges and then transported to the laboratory. The agar material from each device was added to 1 ml of sterilized water including 3.5% artificial sea salt, homogenized with a sterile stick, vortexed, diluted with sterile water including 3.5% artificial sea salt, subcultured on media as described above with 1.5% agar, and incubated at 20°C. After one week, 60 colonies were randomly selected, pure cultured using 1:10 R2A medium and used for further analysis.

### Standard direct plating cultivation

Conventional standard direct plating (SDP) was performed to compare the SDP results with those of the *in situ* cultivation approaches (Fig. 1). Aliquots from sponge samples prepared in the same manner as the DC cultivation were diluted serially and plated directly on the agar media described above. After incubating for one-week at 20°C, 60 colonies were randomly selected and pure-cultured using 1:10 R2A medium for microbial identification.

### Identification of isolates based on 16S rRNA gene sequencing

Taxonomies were identified via sequencing 610- to 712-bp fragments of the 16S rRNA gene. The colony material was used directly as a PCR template. The 16S rRNA gene was amplified using the universal primers, 27F (5’-AGAGTTTGATCCTGGCTCAG-3’) and 1492R (5’- GGTTACCTTGTTACGACTT-3’), with a KOD FX Neo system (Toyobo, Osaka, Japan), then purified using a fast gene purification kit (Nippon Genetics, Tokyo, Japan). The purified PCR products were sequenced with a commercial sequencer (Takara Bio, Shiga, Japan) by fluorescent dye terminator sequencing. The sequences were compared with those available in GenBank (www.ncbi.nlm.nih.gov) using Molecular Evolutionary Genetics Analysis (MEGA software, Tempe, AZ, USA) to determine their closest relatives. Distance matrices and phylogenetic trees based on 16S rRNA sequences were calculated among the isolated OTUs (defined at 97% 16S rRNA gene sequence identity) according to the Kimura two-parameter model and neighbor-joining algorithms using the MEGA program (MEGA software, Tempe, AZ, USA). One thousand fast bootstraps were performed to assign confidence levels to the nodes in the trees.

### DNA extraction and amplicon sequencing targeting the 16S rRNA gene

To compare the cultivated bacterial diversity with the microbial molecular signatures in the host sponge, 16S amplicon sequencing was performed based on the 16S rRNA genes. After transferring the marine sponges to the laboratory, samples were washed three times with DNA-free water and homogenized. Genomic DNA from the homogenized tissue was extracted using a FastDNA spin kit for soil (MP Biomedicals, Irvine, CA, USA) per the manufacturer’s guidelines. The extracted genomic DNA was amplified using the amplicon forward primer and amplicon reverse primer with a Hifi hot start ready mix PCR (Kapa Biosystems, Wilmington, MA, USA), then purified using a fast gene purification kit (Nippon Genetics, Tokyo, Japan). DNA was sequenced by Hokkaido System Science (Sapporo, Japan) using an Illumina MiSeq System (Illumina, San Diego, CA, USA). Each read was assigned taxonomically using Quantitative Insights Into Microbial Ecology (QIIME) software (72).

### Effect of the sponge extract on starvation recovery

The colony formation efficiency ratios between the two culture conditions (media with and without the sponge extract) were calculated for each tested strain to examine the effect of the sponge extract on starvation recovery. First, one strain per OTU (defined at 97% 16S rRNA gene sequence identity) from the DC and SDP cultivations were selected and cultured in 5 ml of 1:10 diluted R2A broth with 3.5% artificial sea salt at 20°C (liquid cultures). Each culture was diluted 100 times and placed at 5°C to inactivate them. After 3 days, the liquid culture serial dilutions were inoculated in triplicate on two types of agar media: 1:10-diluted R2A agar medium with 0.1% (vol/vol) of the sponge extract and the same medium without the sponge extract. The sponge extract added to the medium before autoclaving. After 5 days of incubation at 20°C, the colony number ratios between the two culture conditions (with and without the sponge extract) were measured for each tested strain.

### Effect of the sponge extract on the specific growth rate and carrying capacity

Specific growth rates and carrying capacities under the two culture conditions (medium with and without the sponge extract) were measured and compared for each tested strain to examine the sponge extract’s effect on the growth activity. First, one strain per each OTU from the DC and SDP cultivations were selected and cultured (liquid cultures) in the same manner as described above. Next, 5–20 μl of the microbial cell suspension from each liquid culture was inoculated in triplicate into the two media (1:10-diluted R2A medium with 0.1% [vol/vol] of the sponge extract and the same medium without the sponge extract). The sponge extract added to the medium before autoclaving. The cultures were then incubated at 20°C with shaking at 150 rpm. The optical density (OD) was measured at 600 nm using a spectrophotometer (DR 3900, HACH, Metropolitan, MD, USA). The growth curve for the OD 600 value was fitted using the nonlinear regression function with a logistic model in Sigma plot software (San Jose, CA, USA, Systat Software, Inc.). Specific growth rates, as maximum rates of change, were determined to *μ* by the change in OD 600 on a logarithmic scale as time. The ratio of the specific growth rate (*μ*) between the values measured under the two culture conditions (with and without the sponge extract) was calculated for each tested strain. The carrying capacity (maximum growth) was determined by its maximum value on the fitted growth curve.

## Data availability

Newly determined sequence data have been deposited in GenBank (www.ncbi.nlm.nih.gov) under accession numbers MK672856 to MK672868 for isolates from diffusion chamber, and MK674856 to MK674892 for isolates from standard direct plate cultivation.

## Acknowledgment

This work was supported by JSPS KAKENHI Grant Numbers JP17F17098, JP25630383, JP18K19181. D. J is an International Research Fellow of the Japan Society for the Promotion of Science. We thank Traci Raley, MS, ELS, from Edanz Group for editing a draft of this manuscript. We thank Slava S. Epstein for valuable comments and discussions on our data.

## Supplemental Materials

Table S1. Phylogenetic affiliations of isolates with the DC method on the basis of 16S rRNA gene sequences.

Table S2. Phylogenetic affiliations of isolates with the SDP method on the basis of 16S rRNA gene sequences.

**Figure S1.**
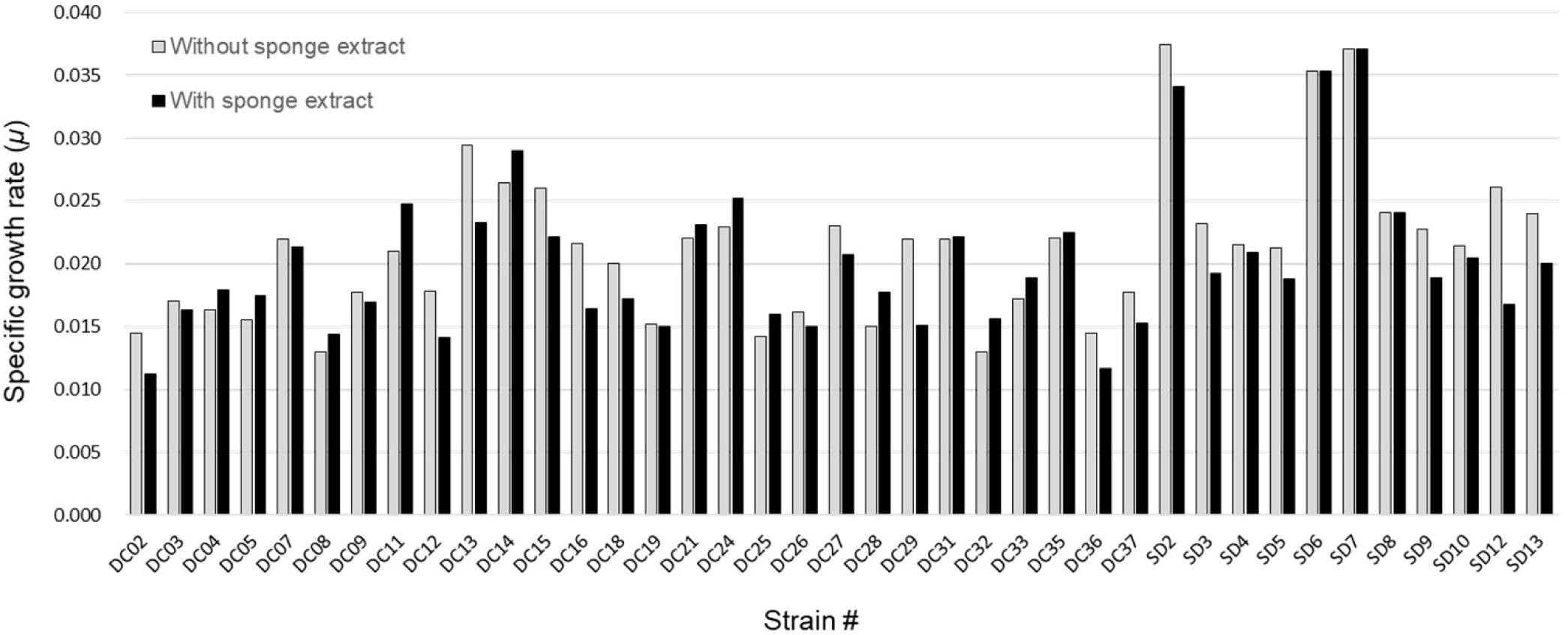
The specific growth rate for each tested strain measured under the two culture conditions (the medium with and without the sponge extract).

